# Chitosan gold nanoparticle-based dot-blot assay for sensitive visual detection of histidine-tagged recombinant proteins

**DOI:** 10.1101/2025.02.07.636955

**Authors:** Sahar Mahmoodi, Mohammad Pourhassan-Moghaddam, Hasan Majdi, Mohammad Jafar Maleki

**Affiliations:** Department of Medical Biotechnology, Faculty of Advanced Medical Sciences, Tabriz University of Medical Sciences, Tabriz, Iran; Department of Medical Nanotechnology, Faculty of Advanced Medical Sciences, Tabriz University of Medical Sciences, Tabriz, Iran; Graduate School of Biomedical Engineering, Faculty of Engineering, University of New South Wales, Sydney, Australia

**Keywords:** Recombinant protein, Chitosan-gold nanoparticles, Dot blots immunoassay, Visual detection, Gold enhancement system

## Abstract

There are various techniques for detecting recombinant proteins such as western blot, SDS-PAGE, ELISA and fluorescence microscopy. However, these methods are difficult to handle, time-consuming and need special tools. We developed a rapid, inexpensive, sensitive, and straightforward approach to address these problems using an Antihistidine biosensor. Colloidal gold nanoparticles (GNPs) were synthesized by chitosan as reducer and stabilizer via the green synthesis method. Then, anti-His tag antibody was immobilized on the Chitosan-gold nanoparticles (CS-GNPs) surface to visually detect spotted target protein on nitrocellulose (NC) membrane. Our results showed that the designed dot-blot immunoassay can detect histidine-tagged recombinant proteins with the limit of detection (LOD) of 1µg/ml without any signal enhancement directly from cell lysate within 5 minutes. In an extra step, we applied HAuCl4 and NH2OH\ HCl as a gold enhancement system and increased the detection sensitivity to 0.1 µg/ml. The results showed that the developed assay can rapidly detect production of recombinant proteins, and it can be used as a screening method in low-resource laboratories.

## 1. Introduction

Recombinant proteins are produced to meet the needs for elucidation of genes’ function, diagnosis and treatment of diseases, and developing novel industrial compounds. The biotechnology methods enable expressing proteins very easily, whereas the development of simple methods for detecting proteins is still challenging. There are various methods for detecting recombinant proteins such as electrophoresis-based assays-Western blot, Sodium Dodecyl Sulphate Poly Acrylamide Electrophoresis (SDS-PAGE), enzyme-linked immunosorbent assay (ELISA) and fluorescence microscopy. However, these methods are difficult to handle, time-consuming, and need special tools [1, 2]. An ideal detection platform is a simple, inexpensive method that rapidly recognizes high sensitivity and specificity targets.

The poly histidine-tag (His-tag) is one of the most famous affinity tags fused to recombinant proteins during cloning for affinity separation. This tag is used as an appropriate tool in the detection and isolation of recombinant proteins. Its size is about 1 kD, so it scarcely affects the structure and function of recombinant proteins. Also, it can easily be linked to a desirable gene and is effectively expressed in various expression systems, comprising E. coli, yeast, insect cells, plant cells and mammalian cells. Consequently, the desirable recombinant protein could be detected by monitoring the His-tag production in host cells [2, 3]. These encouraged us to apply this tag in order to detect recombinant proteins from cell lysates directly.

On the other hand, in recent years, the electrochemical sensor has been highly regarded as an alternative for these methods [4-7]. The electrochemical sensors are economical, easy to fabricate and miniaturize. Moreover, there are various strategies for improving their efficacy and signal enhancement [6-9].

In a study, Gandarilla et al. designed an electrochemical immunosensor to detect Plasmodium falciparum histidine-rich protein 2 antigen (Ag-PfHRP2) as malaria biomarker in humans serum samples. They used dihexadecyl phosphate polymer (DHP) to directly immobilize *Pf*HRP2 antibody on a gold electrode [4]. In another study, a label-free potentiometric immunosensor was developed to detect recombinant human myelin basic protein (rhMBP). They successfully detected rhMBP in the range of 0.10−20.00 μg/mL with a limit of detection of 50.00 ng/mL[5]. Recently, Ren *et al*. designed an electrochemical biosensor by thionine-chitosan-gold nanoparticles (Chit-GNPs) decorated with horseradish peroxidase (HRP) and anti-His tagged protein monoclonal antibody. They obtained a LOD of 3.3 pg/ml (S/N = 3) [9].

Although these biosensors are more rapid than the conventional methods for detecting his-tagged proteins, they have some limitations, such as complex fabrications, high costs, and time-consuming procedures.

Among nanoparticles, gold nanoparticles (GNPs) have long been applied as pivotal tools in biosciences because of their unique optical properties, biocompatibility, controllable size distribution, synthesis flexibility, and visual detection ease. However, the main challenges in using GNPs are surface modification, biofunctionalization and stability of nanoparticles[10].

To address these challenges, we used chitosan to synthesize GNPs, and immobilize antibodies on the nanoparticles. Chitosan is a natural linear polycationic polysaccharide and has special characteristics such as biocompatibility, hydrophilicity, biodegradability and large numbers of reactive hydroxyl and amine groups on its surface for immobilization of biomolecules. This polymer provides the functional groups on GNPs surface and helps stabilize the GNPs colloidal suspension. Various studies applied this natural polymer as a gold nanoparticles coating for different biomedical applications, particularly sensing [11-13].

This research reports on the synthesis of anti-His-tag antibody conjugated CS-GNPs solution (Ab-CS-GNPs) by glutaraldehyde as a green immunonanoprobe to detect recombinant His-tagged proteins from cell lysates visually. In a further step, in order to increase its sensitivity, the signal enhancement system was used based on the GNPs seeding and enlargement. The gold enhancement is a promising method for increasing the detection sensitivity in lateral flow immunoassays or immune-dot blot assay. We mixed HAuCl4 and NH2OH\ HCl to increase the size of immunogold nanoparticles fixed on the nitrocellulose papers [14].

## 2. Materials and Methods

### 2.1. Materials

Gold(III) chloride trihydrate (HAuCl4·3H2O), low molecular weight chitosan (50-190 KDa), phosphate-buffered saline (PBS), bovine serum albumin (BSA), hydroxylamine hydrochloride (NH2OH\ HCl), EDC (1-Ethyl-3-(3 dimethylaminopropyl)-carbodiimide) and NHS (N-hydroxysuccinimide) were obtained from Sigma-Aldrich. His-tag antibody produced in rabbit was purchased from Covalab, France. Tween-20, acetic acid, glutaraldehyde (GA), and nitrocellulose membrane (NC) were purchased from Merck. Other chemicals applied in this study were analytically pure and with maximum quality. Ultrapure deionized water (DI) was used in all experiments.

### 2.2. Method

#### 2.2.1 One-step synthesis of CS-GNPs

In this research, chitosan was used to synthesize stable colloidal gold nanoparticles according to Huang, H. and X. Yang method [15]. A solution of 10 mg/ml chitosan in 1% acetic acid was mixed with 1mmol/l HAuCl4 in 1:1 ratio and incubated at 65°C with shaking. 2 hours after observing the red color in the reaction, the temperature was reduced to 25°C while the mixture was left shaking for overnight. The synthesized colloidal solution was characterized by UV-Vis spectroscopy and TEM after centrifugation and dilution with deionized water. This solution can be stored at 4°C for several months.

#### 2.2.2 Bio-conjugation of antibody with the surface of CS-GNPs

To immobilize antibodies on CS-GNPs surface, we applied Glutaraldehyde (GA) and EDC/NHS. As a bifunctional cross-linking chemical, GA was used to covalently connect amine groups in the chitosan layer on the surface of CS-GNPs to antibodies [16].

Initially, the optical density (OD) of CS-GNPs was reduced to 0.4 (0.73 nM) by dilution in Ultrapure DI water. For the preparation of GA-CS-GNPs, GA (at a final concentration of 1%) was incubated with CS-GNPs for 40 min at 25°C. After this time, the excess GA was washed off by centrifugation. 3ul His-tag antibody (1 mg/ml) was mixed to 100 ul of GA-CS-GNPs in different pH values (5–9). After five minutes, 20 ul of 10% NaCl was mixed with each of them. After two hours, the minimum pH in which the red color of the solution did not change was selected as the optimum pH. Next, the antibody solution from 2 to 20 ul (0.1 mg/ml) was added to the microtubes containing 100 ul of fresh colloidal gold solution with the optimum pH. In the following, 20 ul of 10% NaCl was added to the microtubes. After 2 hours, the least amount of antibody in which the color of solution did not change was considered the optimum amount of antibody for stabilizing the gold nanoparticles. Accordingly, a proper amount of His-tag antibody was added dropwise to GA-CS-GNPs with optimum pH and incubated at 4 °C for overnight. Then, the mixture was washed with PBS by centrifugation at 10000 rpm for 1 hour. After washing steps, for blocking the non-specific binding sites on the surfaces of the particles, the solution was incubated with BSA (5%) in PBS (0.01 M, pH7.4) for 20 min at 25°C. Eventually, the conjugate was washed with PBS twice and suspended in PBS again for storing at 4°C.

We adopted Narayan *et al*. method [17] for conjugation by EDC/NHS. In brief, 1 mL of antibody solution (1 mg/ml) was mixed with 1.5 mg EDC and 1.1 mg NHS. Next, various concentrations of this solution (0.01, 0.1 and 1 mg/ml) was prepared. Then, 0.3 ml from each prepared concentration was mixed with the 0.7 ml CS-GNPs solution and incubated at 4°C for overnight. For blocking the non-specific binding sites on the particle’s surface, the solution was incubated with BSA (5%) at 4^°^c for 45 min. Finally, to remove excess BSA, the solution was pelleted and re-suspended in PBS.

#### 2.2.3 Characterization of CS-GNPs and antibody-CS-GNPs

UV/Visible double beam spectrophotometer (CECIL 7500 CE, UK) was applied to record absorbance spectra. The wavelength range was 400–800 nm. Zeta potential of the GNPs and size based on z-average were determined using photo-correlation spectroscopy (PCS) (Zetasizer-ZS, Malvern, UK). Scanning electron microscopy (SEM), (SEM; 70 EM3200, KYKY Instruments, China) and Transmission electron microscopy (LEO 906, Zeiss-Leica Cooperation, Germany) were used to report the size and morphology of as-prepared CS-GNPs. Digimizer software was applied for the determination of the average diameters of nanoparticles in electron microscopy images. Before use, all glassware was washed using deionized water and sonicated 30 min in ultrasonic bath (Witeg Labortechnik GmbH, Germany).

#### 2.2.4 Visual detection of His-tagged protein

In order to study the selectivity of the as-prepared His-tag antibody-CS-GNPs, His-tagged protein, mouse IgG and goat IgG were spotted (2µl of 1mg/ml solution of each) on the nitrocellulose papers and dried at 25^°^C for 10 min. Subsequently, papers were immersed in 3% BSA solution and rinsed with PBS. Then, the papers were incubated in His-tag antibody-CS-GNPs (Ab-CS-GNPs) solution for 5 min, rinsed with PBS, and dried at 25^°^C. The capability of the assay in the direct detection of his-tagged protein from cell lysate was also tested. Lysate of induced E.coli cells was used as the sample in the dot blot assay.

Next, the assay’s limit of detection was determined: we used Ab-CS-GNPs solution to identify different concentrations of (100, 10, 1, and 0.1 µg/ml) target protein. In order to reduce the limit of detection, the signal enhancement system was used based on gold enlargement using HAuCl4 and NH2OH\HCl. The nitrocellulose papers were immersed in the mixture of 100 μl of HAuCl4 (1 mM) and 500 μL NH2OH\ HCl (200 mM) for 2-3 min at 25°C and then washed with water[18].

## 3. Results and Discussion

Bright red colour CS-GNP solution was successfully produced using chitosan as a reducer and stabilizer through a green synthesis method (Fig. 1A) with LSPR band at 523 nm that suggested the formation of spherical CS-GNPs of size < 25 nm (Fig. 1B). DLS analysis showed an average size of 50 nm with PDI of 0.226 and a zeta-potential of +34 mV (Table 1), indicating the narrow size and stability of CS-GNPs. SEM and TEM revealed dispersed spherical particles of 40 nm (Fig.2 B, D) and 17 nm (Fig. 2 A, C), respectively.

**Fig. 1.**
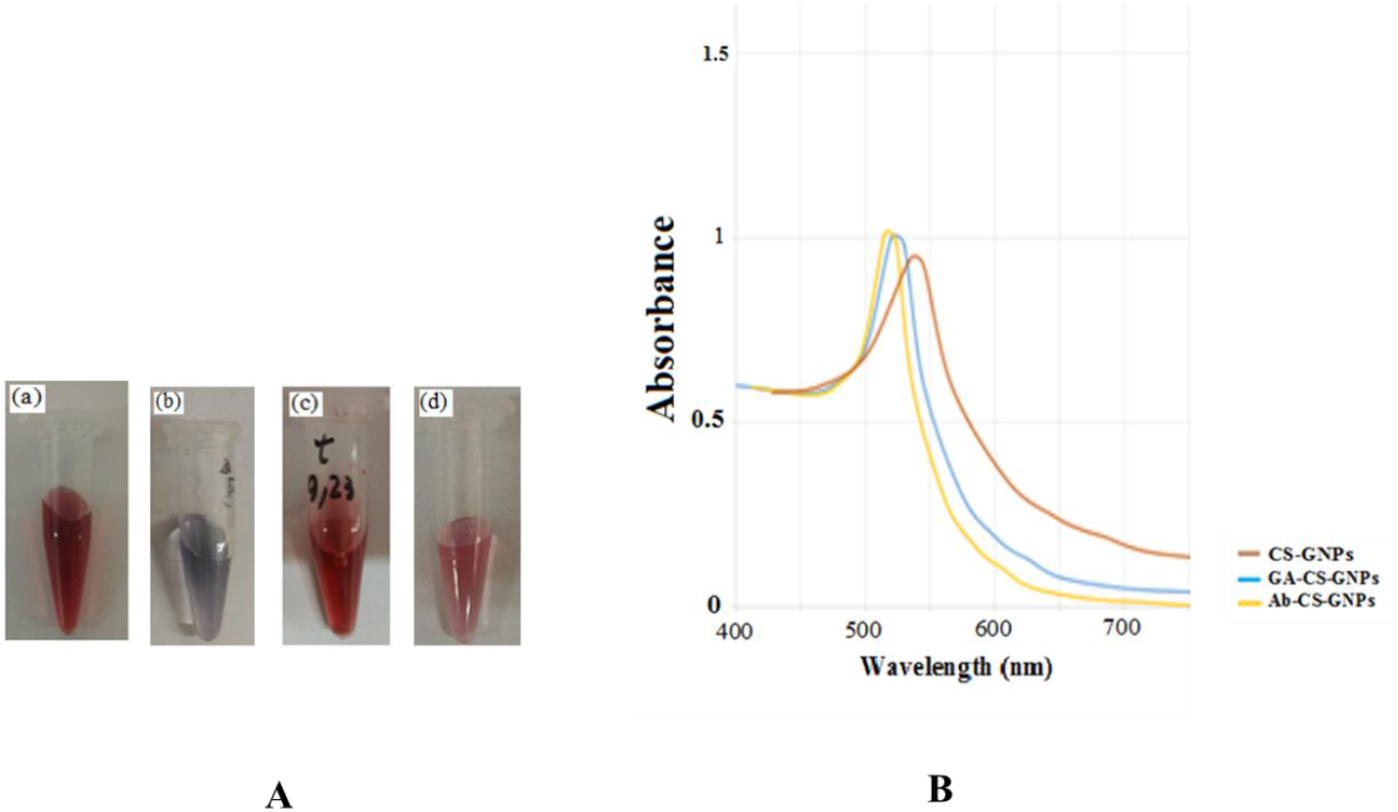
**(A)**ctures of Chitosan-gold solution during the Ab conjugation Process by Glutaraldehyde and EDC/NHS solution ; (a) CS-GNPs, (b) CS-GNPs after addition of EDC/NHS solution,(c) CS-GNPs after addition of GA solution (GA-CS-GNPs), (d) GA-CS-GNPs after addition of Ab and washing with PBS. **(B)** UV–Vis absorbance spectrum of CS-GNPs during the Ab conjugation by glutaraldehyde as a linker.

**Table 1.** Hydrodynamic average diameter, PDI and Zeta Potential of CS-GNPs, GA-CS-GNPs and Ab-CS-GNPs (p < 0.05). Information was presented as mean ± SD.

| **Sample**<br>**(mv)** | **Average diameter (nm)** | **PDI** | **Zeta Potential** |
| --- | --- | --- | --- |
| **CS-GNPs** | 50 ± 4.08 | 0.226 ± 0.07 | + 34 ± 7.6 |
| **GA-CS-GNPs** | 88 ± 6.80 | 0.245 ± 0.03 | +24 ± 3.6 |
| **Ab-CS-GNPs** | 112 ± 7.01 | 0.287 ± 0.05 | -22 ± 4.02 |

**Fig. 2.**
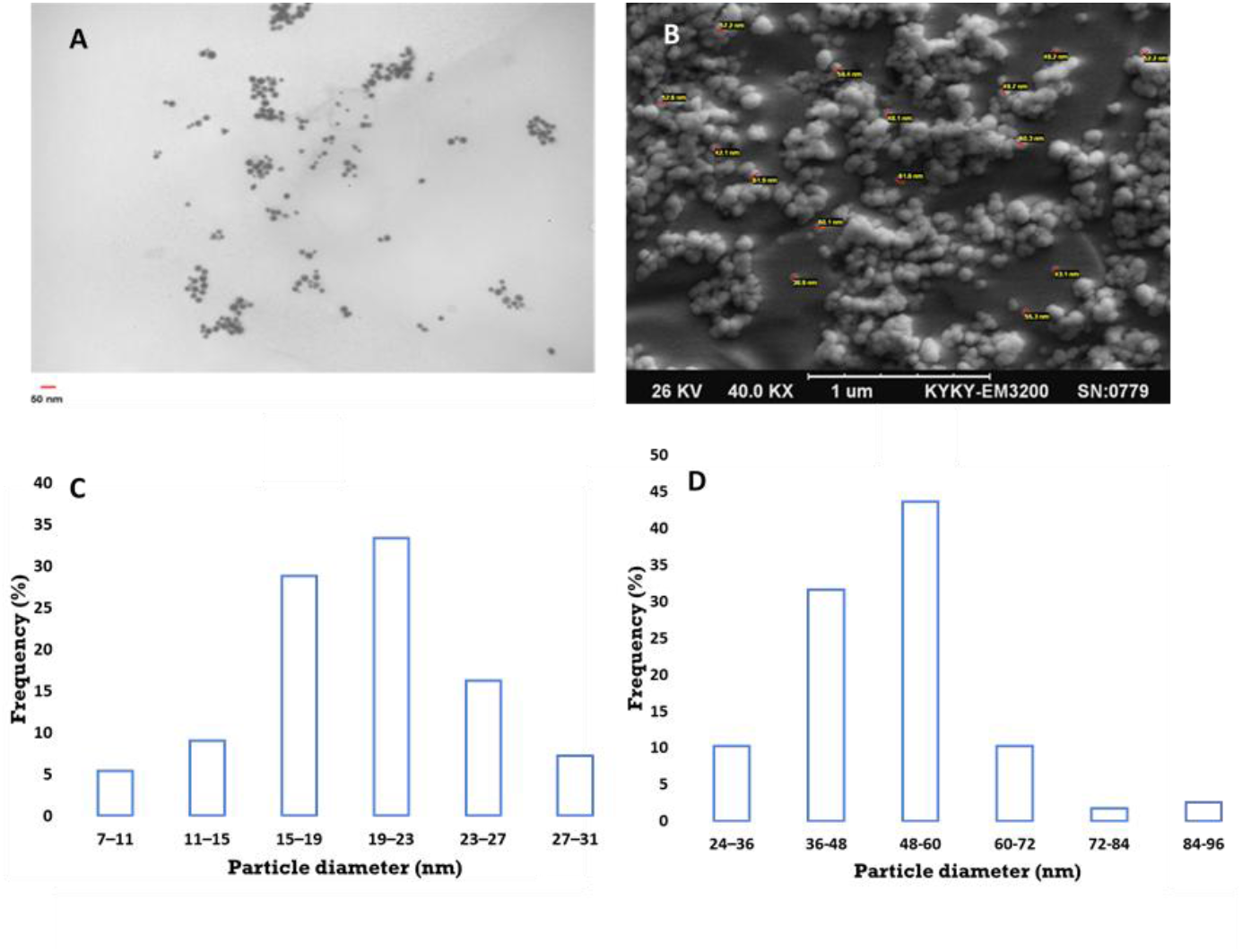
(A) TEM image (B) SEM image (C) TEM size distribution histogram and (D) SEM size distribution histogram of Chitosan-gold nanoparticles.

The greater size of CS-GNP reported by SEM, reflects the formation of a thick chitosan layer around the metallic cores of CS-GNPs [19].

On the other hand, the formation of pseudo-clusters and water layers on CS-GNPs had resulted in the difference between the DLS and the SEM measurements [20].

We used EDC/NHS and GA solution to conjugate antibodies on CS-GNPs surface (Fig.1A;a). During the conjugation process, the addition of EDC/NHS solution aggregated CS-GNPs (Fig.1A;b). On the contrary, GA created more stability in the solution (Fig.1A;c,d). This stability can be attributed to cross-linking of the amine groups of chitosan layers of the adjacent CS-GNPs and the formation of the cross-linked CS-GNPs [21]. According to these results, we selected GA for the conjugation of Ab on the surface of CS-GNPs. To obtain the optimum amount of Ab for the conjugation process, we prepared various concentrations of the anti-his tag antibody solution consisting of 0.001, 0.01 and 0.1 mg/ml. As the concentration of antibody increased from 0.001 to 0.1, the stability of the particles increased. The particles aggregation in low concentration of Ab (0.001 mg/ml) could be because of the slight variations in surface charge of CS-GNPs [22]. After the conjugation step, the UV/Vis spectra of CS-GNPs showed a right shift from 527 nm to 531 nm (Fig.1B), indicating immobilization of anit-His-tag antibody on the surface of CS-GNP, and formation of Ab-CS-GNPs [23, 24]. DLS was used to monitor changes in size and zeta-potential of CS-GNPs during the conjugation process. The activated CS-GNPs showed a size of 88 nM and a zeta-potential of +24 mV. Incubation of the activated particles with anti-his-tag antibody increased particles’ size to 112 nm. Furthermore, it resulted in negative zeta potential of - 22 mV, suggesting immobilization of the antibody molecules on CS-GNP surfaces which modified the surface characteristics of CS-GNPs after conjugation (Table 1) [25]. In conclusion, the optimum pH for conjugation of anti-his tag antibody with CS-GNPs was obtained 7.6. Also, at this pH, the minimum amount of antibody was determined 0.01 mg/ml to stabilize the GA-CS-GNPs.

After preparing Ab-CS-GNPs, a dot-blot assay was used for visual and rapid detection of his-tagged protein samples. A red spot was observed only where target protein was spotted on the membrane, proving the specific binding of Ab-CS-GNPs with the target protein (Fig.3). This result demonstrates that the designed conjugate has enough detection specificity. It is worth mentioning that we could detect target protein from induced E. coli cell lysates. When cell lysate was spotted on the nitrocellulose membrane, red color was visible even in the presence of a mixture of proteins in the cell lysate (Fig.4). For the next experiment, we used Ab-CS-GNPs solution to identify different concentrations of (100, 10, 1, and 0.1 µg/ml) target protein (Fig. 5). The limit of detection (LOD) was obtained 1 µg/ml without any signal enhancement. This result proves that the present visualization technique is highly sensitive and applicable as a reliable technique where instruments are not accessible. In order to improve the sensitivity of the assay, a signal enhancement step was included based on gold enlargement using HAuCl4 and NH2OH\HCl [14, 18]. Thermodynamically, hydroxylamine (NH2OH) reduces Au^3+^ ions. In addition, Au^0^ surfaces accelerate this reaction. As a result, the added Au3+ ions were consumed to form larger particles [14]. This process results in color changes that can be visualized by the naked eye. Our results demonstrated that the detection sensitivity was considerably raised to 0.1μ g/ml (picomole level) (Fig.6).

**Fig. 3.**
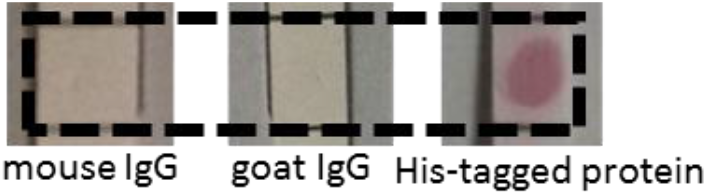
Recognition of His-tagged protein by His-tag antibody-CS-GNPs (red color formation), no color was formed for mouse IgG and goat IgG.

**Fig. 4.**
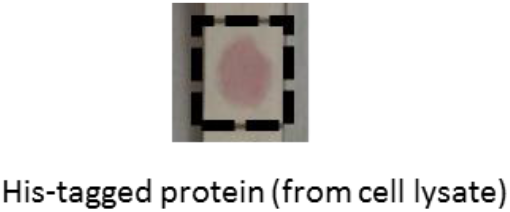
Recognition of His-tagged protein by His-tag antibody-CS-GNPs from cell lysate

**Fig. 5.**
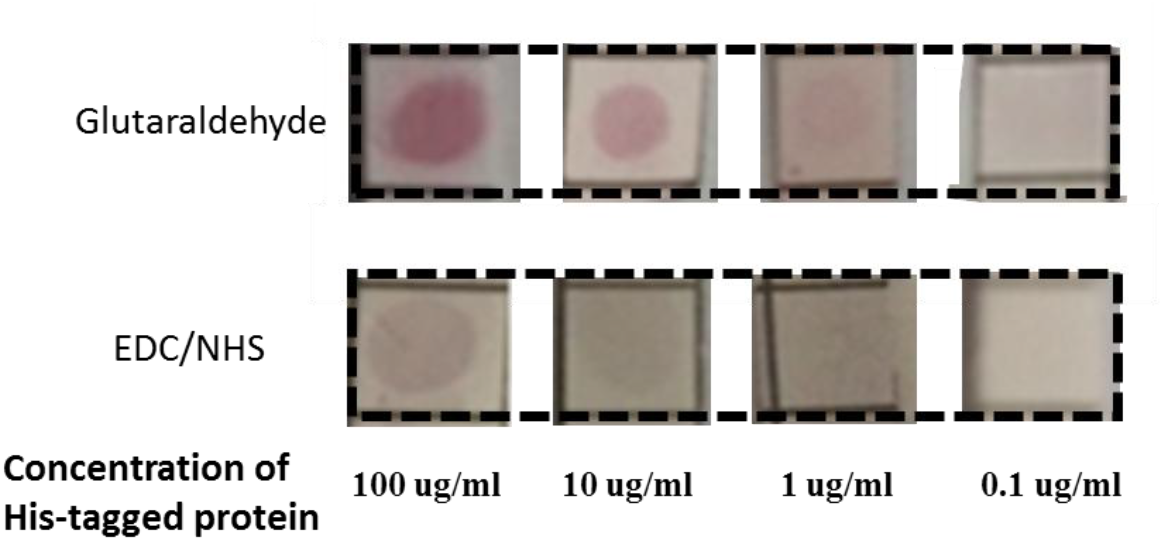
Intensity of red spots formed with different concentration of His-tagged protein (100µg/ml, 10 µg/ml, 1 µg/ml, 0.1 µg/ml), intensity of the red spot decreases from left to right.

**Fig. 6.**
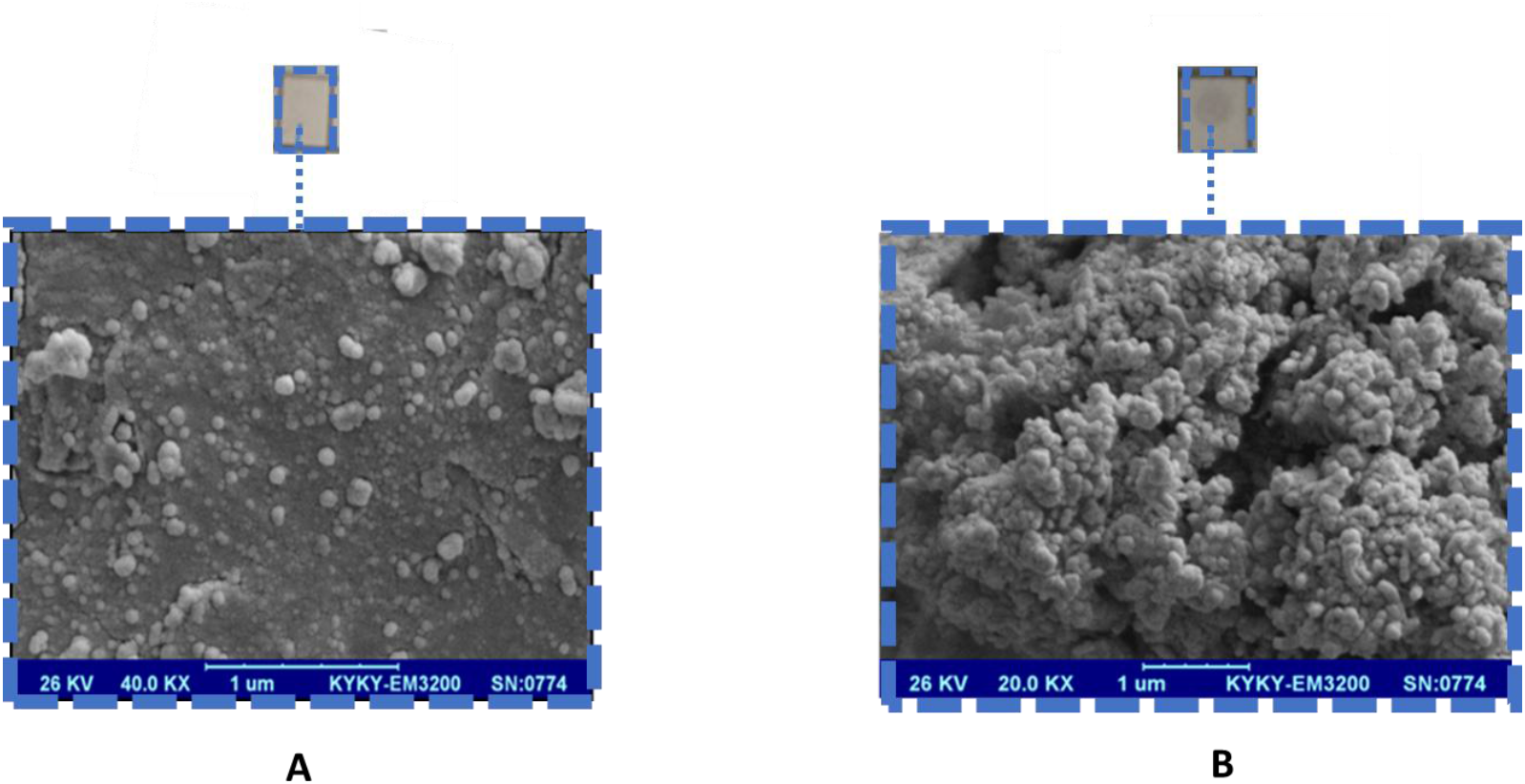
SEM images and intensity of red spot formed with concentration of 0.1 µg/ml from His-tagged protein before (A) and after (B) signal amplification by gold enhancement with HAuCl4 and NH2OH\ HCl.

## 4. Conclusion

In this research, we have designed a novel green immunonanoprobe to directly detect recombinant His-tagged protein from cell lysates at low concentration by the naked eyes. We immobilized antibodies directly without ligand exchange on the Cs-GNPs surface by covalent bonding. The limit of detection was obtained 1 µg/ml. We could significantly increase the detection sensitivity to 0.1μg/ml using a gold enlargement system. The advantages of this approach consist of rapid, low cost, widespread applicability, eco-friendly, simplicity, high stability, high specificity, high sensitivity and functionality for a long period of time [7, 26].

## 5. Acknowledgement

This article has been derived from the MSc thesis number “95/2-2/1”. The authors would like to thank the Research Deputy of Tabriz University of Medical Sciences for their financial support.

## Notes

### Competing Interest Statement

The authors have declared no competing interest.

